# Early life adversities affect expected value signaling in the adult brain

**DOI:** 10.1101/2023.06.19.545539

**Authors:** Seda Sacu, Magda Dubois, Pascal-M. Aggensteiner, Maximilian Monninger, Daniel Brandeis, Tobias Banaschewski, Tobias U. Hauser, Nathalie Holz

**Author notes:** Corresponding Author: Dr. Nathalie Holz, Donders Centre for Cognitive Neuroimaging, Donders Institute for Brain, Cognition and Behaviour, Radboud University, Kapittelweg 29, 6525 EN Nijmegen, Department of Child and Adolescent Psychiatry and Psychotherapy, Central Institute of Mental Health, Medical Faculty Mannheim / Heidelberg University, J 5, 68159 Mannheim, Germany, Tel: +49 621 /1703-4904. These authors contributed equally (Seda Sacu, Magda Dubois, Tobias U. Hauser, Nathalie Holz).

## Abstract

**Background:** Early adverse experiences are assumed to affect fundamental processes of reward learning and decision-making. However, computational neuroimaging studies investigating these circuits are sparse and limited to studies that investigated adversities retrospectively in adolescent samples.

**Methods:** We used prospective data from a longitudinal birth cohort study (n=156, 87 females, mean age=32.2) to investigate neurocomputational components underlying reinforcement learning in an fMRI-based passive avoidance task. We applied a principal component analysis to capture common variation across seven prenatal and postnatal adversity measures. The resulting adversity factors (factor 1: postnatal psychosocial adversities and prenatal maternal smoking, factor 2: prenatal maternal stress and obstetric adversity, and factor 3: lower maternal stimulation) and single adversity measures were then linked to computational markers of reward learning (i.e. expected value, prediction errors) in the core reward network.

**Results:** Using the adversity factors, we found that adversities were linked to lower expected value representation in striatum, ventromedial prefrontal cortex (vmPFC) and anterior cingulate cortex (ACC). Expected value encoding in vmPFC further mediated the relationship between adversities and psychopathology. In terms of specific adversity effects, we found that obstetric adversity was associated with lower prediction error signaling in the vmPFC and ACC, whereas lower maternal stimulation was related to lower expected value encoding in the striatum, vmPFC, and ACC.

**Conclusions:** Our results suggested that adverse experiences have a long-term disruptive effect on reward learning in several important reward-related brain regions, which can be associated with non-optimal decision-making and thereby increase the vulnerability of developing psychopathology.

## Introduction

Being able to adapt and learn about one’s environment is critical for successfully navigating the world (1). Developing accurate predictions about future events and updating them based on novel information becomes especially important in dynamic environments where constant change is present (2). However, these fundamental processes of feedback learning have been found to be impaired across a range of mental disorders (3,4). Early adverse environments are also believed to alter reinforcement learning processes as inconsistencies in feedback contingencies (5) and suboptimal conditions for neurocognitive development (6) are prevalent in adverse rearing environments.

Expected value (EV) and prediction error (PE) are two important interrelated processes that underlie successful reinforcement learning (7). PE occurs when there is a discrepancy between expected and actual outcomes, and serves as a teaching signal by allowing the organism to update the EV of a future event (8). At the neural level, several brain regions were found to involve in EV and PE signaling including the striatum (7–9), ventromedial prefrontal cortex/medial orbitofrontal cortex (7,10), anterior cingulate cortex (11), and amygdala (1,9). Lower EV and PE signaling in these regions has also been identified in several psychiatric conditions (12–15).

Several neuroimaging studies have reported a relationship between adverse experiences and alterations in the reward circuitry (16–20), however, research investigating this using computational neuroimaging approaches remains scarce. Computational neuroimaging brings new insights by taking into account other important information regarding the stimulus (e.g., probability and magnitude), which allows modeling the cognitive process beyond the simple stimulus-response relationship (10). To date, only a few previous studies investigated the association between early adverse experiences and EV/PE signaling using a model-based fMRI, suggesting that adversities may indeed affect EV and PE signaling in the adolescent brain (21–23). However, these studies included only single measures of adversity (21–23), despite the fact that adversities tend to co-occur and accumulate over time (24). Therefore, a comprehensive approach is needed to study the long-term effects of developmental risks on EV and PE signaling in the adult brain. Moreover, although previous evidence exists that prenatal and perinatal adversities have an impact on child behavior and brain development (25), their relations to the reward system have barely been studied (26) and need to be explored.

Here, we aimed to investigate the effect of a lifespan adversity profile on EV/PE signaling in a cohort of adults followed since birth using a reinforcement learning paradigm. Risk measures were collected across the development and included prenatal, perinatal, and postnatal factors. Based on previous research (21–23), we hypothesized that adverse experiences across the development would be associated with lower EV and PE encoding in the striatum, ventromedial prefrontal cortex (vmPFC), and anterior cingulate cortex (ACC). Furthermore, lower EV and PE encoding in these regions would be associated with higher psychopathology symptoms in adulthood.

## Methods and Materials

### Participants

The present study was conducted within the framework of the Mannheim Study of Children at Risk, which is an ongoing longitudinal birth cohort study. The initial sample included 384 children born between 1986 and 1988. The infants were recruited from two obstetric and six childreńs hospitals in the Rhine-Neckar region of Germany. The participants were followed from their birth up to around the age of 33 years (age range: 31.7-34.5 years) across 11 assessment waves.

At the last assessment wave, 256 participants (67%) agreed to participate in the study and completed psychological measurements. fMRI data for the passive avoidance task was available for 170 participants. After the quality check (Supplementary Material S1 for exclusion criteria), the sample size was reduced to 156 participants (Table 1). At the time of the fMRI assessment, 22 (14%) participants had current psychopathology including major depressive disorder (n=7), anxiety disorder (n=9) and alcohol and substance abuse (n=5) and attention deficit hyperactivity disorder (n=1), which was assessed using the German version of the Structured Clinical Interview for DSM-IV (27). The study was approved by the ethics committee of the University of Heidelberg. All participants gave informed consent and were financially compensated for their contribution.

**Table 1.** Sample Characteristics.

|  | N=156 |
| --- | --- |
| Age, M(SD) | 32.4(0.4) |
| Sex, N, F/M | 87/69 |
| Maternal Smoking, N, non-/moderate/heavy smoker | 116/17/23 |
| Maternal Stress, M(SD), range | 2.8(1.9), 0-8 |
| Obstetric Adversity, N, no/moderate/high risk | 62/83/11 |
| Maternal Stimulation <sup>a</sup>, M(SD), range | 0.31(2.4), 6-7.2 |
| CTQ total, Median(IQR), range | 28(6), 25-87 |
| Family Adversity, M(SD), range | 3.5 (2.3), 0-10 |
| Stressful Life Events <sup>b</sup>, M(SD), range | 0(6), -11.2-22.2 |
| Internalizing Symptoms, Median(IQR), range | 5(8), 0-40 |
| Externalizing Symptoms, Median(IQR), range | 7(12), 0-45 |
| ADHD, Median(IQR), range | 4(6),0-24 |
| Antisocial Personality, Median(IQR), range | 2(3),0-19 |
| Anxiety, Median(IQR), range | 3(4),0-9 |
| Avoidant Personality, Median(IQR), range | 1(4), 0-11 |
| Depression, Median(IQR), range | 2(5), 0-19 |
| Somatic Problems, Median(IQR), range | 1(2), 0-11 |

<sup>a</sup> We used reversely-coded z-transformed scores. Higher scores indicated lower maternal stimulation.
<sup>b</sup> We used the sum score of z-transformed total scores across the 11 assessment waves.

### Psychological Measurements

#### Lifespan Adversity

The Mannheim Study of Children at Risk was designed to investigate long-term outcomes of psychosocial and biological risk factors on development (28). Consequently, several risk measures were collected across the development (Figure 1). All developmental risk measures were carefully selected based on their impact on psychosocial and psychopathological development (16,17,21,25,26,29,30). For the prenatal period, we included maternal stress (30) and maternal smoking (31), which were measured using a standardized interview during the 3-month assessment. Maternal stress measure contained 11 questions covering negative experiences during pregnancy (e.g., ‘Did you have mood swings/ a depressed mode?’), whereas maternal smoking measured daily cigarette consumption of mothers (1= no, 2= up to 5 per day, 3= more than 5 per day). Obstetric adversity included obstetric complications (e.g., low birth weight, preterm birth, medical complications) as a measure of perinatal risk (28). Postnatal measures included several psychosocial measures. Maternal stimulation was based on video recordings of mother-infant interactions in a play and nurse setting at the 3-month assessment (32), where trained raters evaluated mothers’ attempts (vocal, facial or motor) to draw infants’ attention. We prospectively assessed psychosocial adversities covering family adversity such as parental psychopathology, marital discord from birth to 11 years (28), and stressful life events over the lifespan using the Munich Event List (33). At the age of 23 years, participants filled childhood trauma questionnaire (34). Detailed descriptions for each adversity measure can be found in Supplementary Material S2.

**Figure 1.**
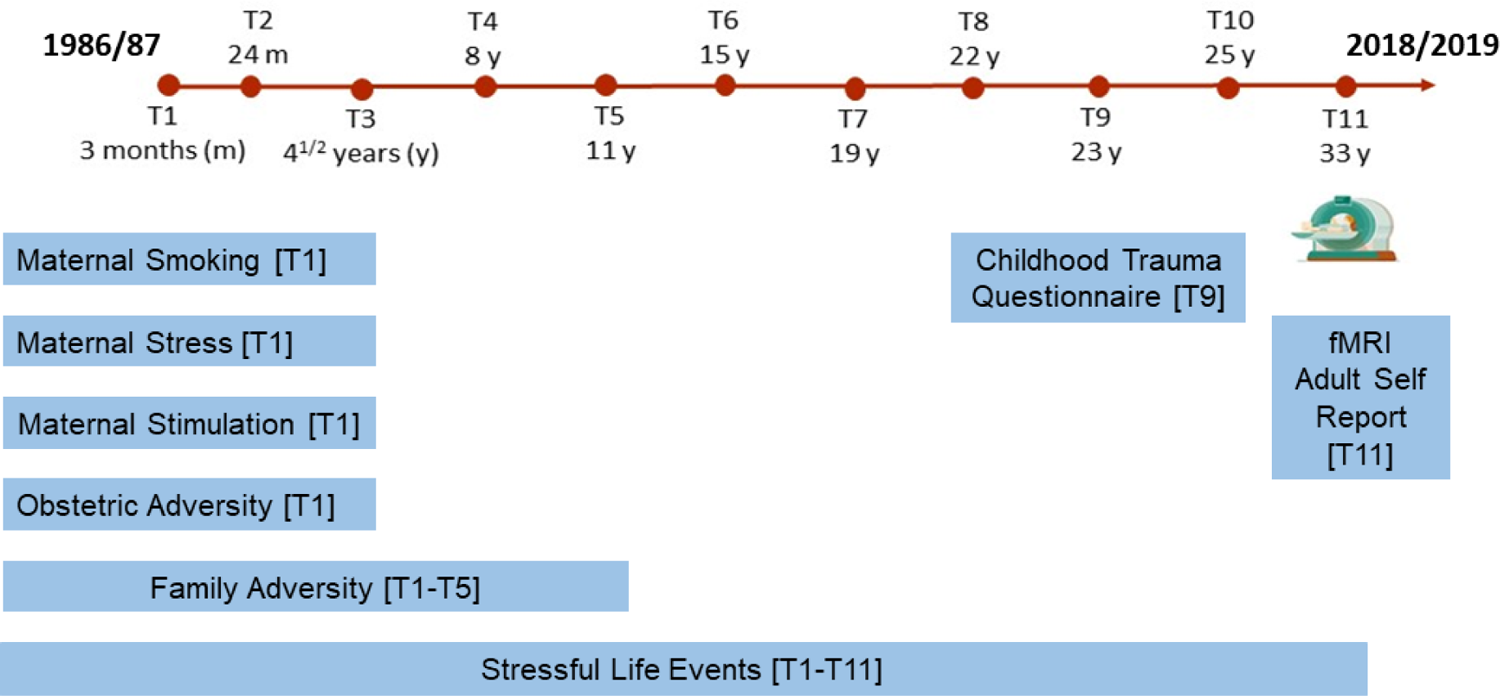
Design of Mannheim Study of Children at Risk.

To reduce the dimensionality while also accounting for their interrelatedness of the adversity measures (Table S1), we applied principal component analysis using the above-mentioned adversity measures (35–37). We identified three components with an eigenvalue > 1, which in total explained 66.8% of the variance in the data (See details in Results).

#### Psychopathology

We used the Adult Self Report (38) to assess current symptoms of psychopathology. The Adult Self Report (ASR) includes summary measurements of psychopathology such as internalizing and externalizing problems. We here used internalizing and externalizing problems total scores to see if general psychopathology scores are associated with adversity factors and brain responses. If we identified an association, we further explored if a specific subscale contributed to this association using the six DSM-oriented ASR subscales including depression, anxiety, avoidant personality, somatic problems, attention deficit and hyperactivity disorder (ADHD) and antisocial personality scales. Due to the high number of testing, we applied a Bonferoni correction (p < 0.05/6 = 0.008) to correct for multiple testing problem.

#### Functional MRI Paradigm

We used a passive avoidance task (14) to measure neural correlates of EV and PE signaling. Each trial started with a presentation (1500 ms) of one of the four colored shapes (Figure 2). During this period, participants had to decide whether to respond or not respond to a shape. A randomly jittered fixation cross (0-4000 ms) followed the presentation of the shapes. If responded, participants received one of the four outcomes: winning 1 €, winning 5 €, losing 1 €, or losing 5 €. Each shape could engender each of these outcomes. However, the feedback was probabilistic. That is, one shape overall resulted in a high reward, one in low reward, one in low punishment, and one in high punishment. If not responded, participants received no feedback. Instead, a fixation cross was presented. Another randomly jittered fixation cross (0–4000 ms) was displayed at the end of each trial. Participants completed 112 trials over two runs.

**Figure 2.**
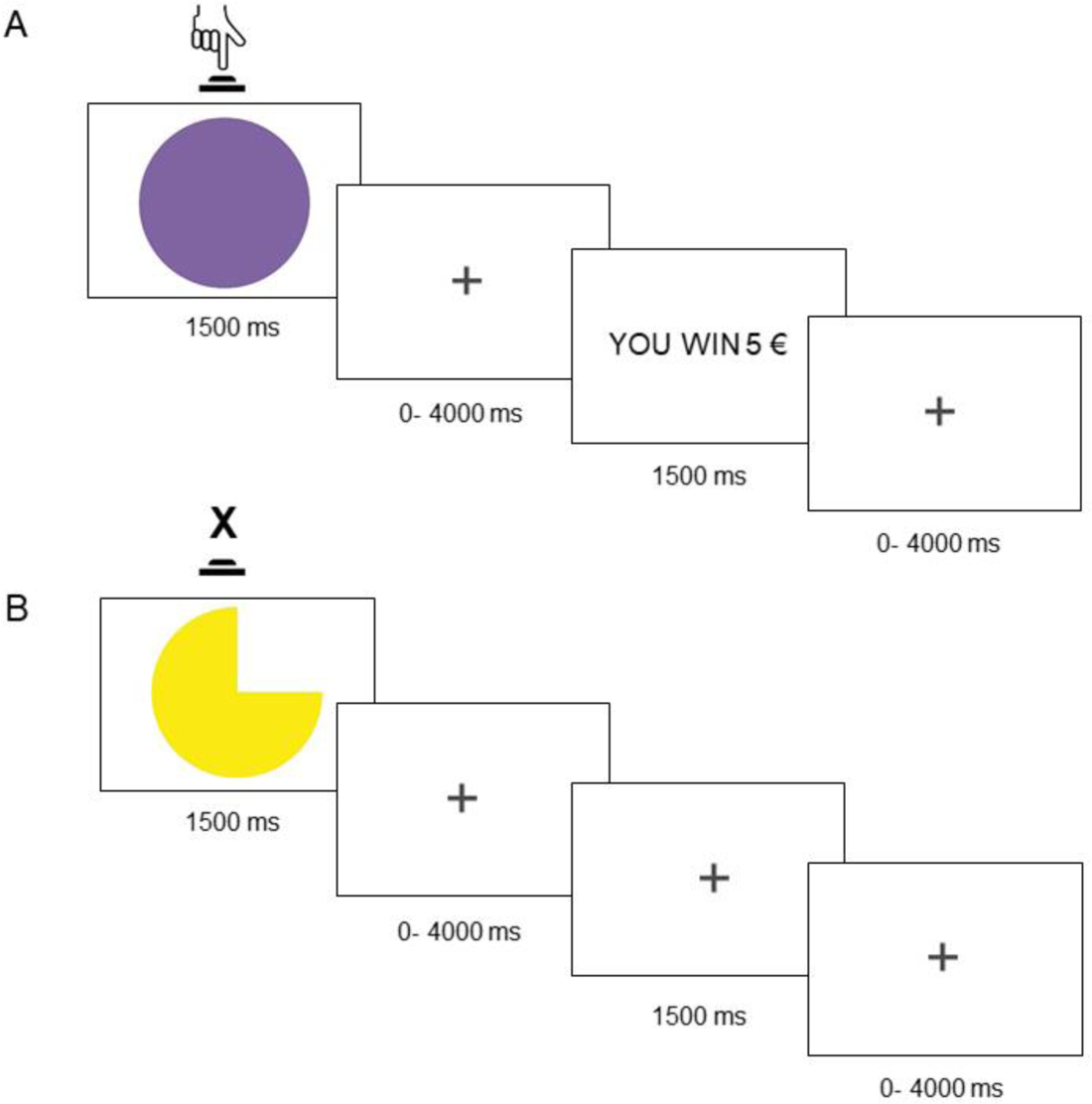
Passive avoidance task. (A) The participant responds to a shape and receives feedback. (B) The participant avoids responding and receives no feedback.

### MRI Data Acquisition and Preprocessing

The functional and structural images were acquired on a Siemens Magnetom Prisma Fit (Siemens, Erlangen, Germany) 3T MRI scanner with a standard 32-channel head coil. During the passive avoidance task, 175 volumes were obtained for each run using a gradient echo-planar sequence sensitive to blood oxygen level-dependent (BOLD) contrast (36 slices, TE= 35 ms, TR = 2100 ms, voxel size = 3 × 3 × 3 mm). Functional data was preprocessed using SPM 12 (https://www.fil.ion.ucl.ac.uk/spm/software/spm12/) applying standard preprocessing steps (Supplementary Material S3).

### Computational Modelling

To understand the observed behavior of participants during decision-making, we compared five Rescorla-Wagner model variations. Each model included three parameters, V0 (i.e., expected value for the first trial), α (learning rate) and β (i.e., inverse temperature), however, they differed in terms of extended SoftMax function parameter (e.g., π, pressing bias) and learning rate (common versus separate learning rates for positive and negative PEs) (39). The best model was determined using Akaike and Bayesian Information Criterion. The winning model was the model with four free parameters (V0, α, β, and π; See Supplementary Material S4 for a detailed description of the models and model comparison procedure).

On each trial t, we calculated PE using the following formula.

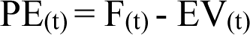

In this formula, the PE for the current trial equals the feedback value (F) minus the EV for the current trial. For the first trial, EV was set to subject-specific parameter estimation obtained from the winning model and was then updated with the following formula.

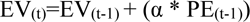

In this formula, the EV for the current trial equals the EV for the previous trial plus the PE for the previous trial multiplied by the learning rate. The learning rate was set to the subject-specific parameter estimation obtained from the winning model. These parameters were then used for the fMRI analysis.

### fMRI Data Analysis

At the first level, we added two onset regressors (at cue and feedback phases) and their parametric modulators (EV and PE, respectively), which were convolved with the hemodynamic response using generalized linear modelling implemented in SPM 12. Six motion parameters were included as covariates of no interest to reduce the motion-related artefacts. At the second level, we performed a one-sample t-test to identify neural correlates of EV and PE signaling. Multiple regression analysis was performed to investigate the associations between three adversity factors and EV and PE signaling in preselected regions of interest: striatum, vmPFC and ACC. These regions reliably take part in EV/PE signaling (7) and show abnormalities in individuals exposed to adverse experiences (21–23). Sex and current psychopathology were included as covariates of no interest. The same regression analysis was also conducted for each adversity measure separately.

Similar to previous reinforcement learning studies (40,41), we thresholded the results with z > 2.3 (cluster-forming threshold) at p < 0.05 (family-wise error corrected (FWE) at cluster level). To correct identified clusters for multiple comparisons, we applied Bonferroni correction to FWE-corrected p values (p < 0.05/3=0.017 for adversity factors and p < 0.05/7=0.007 for single adversity measures). Since our cluster-forming threshold was liberal (41,42), we further performed bootstrapping analysis with 5000 iterations and a 95% confidence interval in IBM SPSS (Version 27) to show the robustness of our results. For that purpose, we extracted mean activation from each significant cluster using the MarsBar toolbox (https://marsbar-toolbox.github.io/).

### Exploratory Analyses

Recently, more attention has been allocated to sensitive period for neural systems, which represents a time window of increased vulnerability to stress (43), and may provide important implications to decide the timing of preventative strategies. To explore the existence of a sensitivity period in which stress exerts enduring effects on reward-related brain activity, we conducted several multiple regression analyses using prospectively collected psychosocial adversity measures. In total, we performed five tests for family adversity (T1-T5) and eleven tests for stressful life events (T1-T11). Since several previous studies have shown that current stress has an impact on neural EV and PE encoding (44–46), we used stressful life events in the last 12 months (T11) to investigate the effect of current stress. Due to the high number of testing and correlative nature, the findings should be considered preliminary.

### Brain-Behavior Association

To investigate the brain-behavior relationship, we extracted mean activation from the clusters significantly related to adversity using the MarsBar toolbox. We used regions-of-interest masks for the extraction if a significant cluster contained several regions-of-interest such as striatum subdivisions and vmPFC to examine the region-specific effect. To this end, putamen, caudate and nucleus accumbens masks were derived from the Melbourne Subcortex Atlas (47) and the vmPFC mask was chosen from a previous study (48). Due to non-normally distributed data, Spearman’s correlation test was performed to identify associations between psychopathology and adversity factors. Furthermore, if we identified association between an adversity factor and psychopathology measure, we performed mediation analysis using the PROCESS toolbox implemented in IBM SPSS (Version 27) to see whether this association was mediated by EV and PE signaling in the brain.

## Results

### Behavioral Results

Principal component analysis identified three adversity factors. The first adversity factor was strongly informed by stressful life events, family adversity, maternal smoking, and childhood trauma questionnaire. The second adversity factor was strongly related to obstetric adversity and maternal stress. The third adversity factor mostly reflected maternal stimulation (Table 2).

**Table 2.** The rotated component matrix for three-factor solution.

|  | Factor 1 | Factor 2 | Factor 3 |
| --- | --- | --- | --- |
| Maternal Stress | 0.26 | **0.76** | -0.02 |
| Maternal Smoking | **0.72** | 0.04 | -0.38 |
| Maternal Stimulation | 0.16 | 0.02 | **0.88** |
| Obstetric Adversity | -0.14 | **0.83** | 0.03 |
| Family Adversity | **0.80** | 0.10 | 0.13 |
| Childhood Trauma Questionnaire | **0.59** | 0.01 | 0.35 |
| Stressful Life Events | **0.83** | 0.02 | 0.19 |

The first adversity factor was associated with higher internalizing (r=0.35, p<0.001) and externalizing (r=0.39, p<0.001) symptoms, specifically to depression (r=0.43, p<0.001), anxiety (r=0.34, p<0.001), avoidant personality (r=0.23, p = 0.005), ADHD (r=0.29, p<0.001), antisocial personality (r=0.32, p<0.001) and somatic problems (r=0.16, p= 0.045). All survived the Bonferroni correction except for the last association. Similarly, the third adversity factor was associated with higher internalizing (r=0.19, p = 0.006) and externalizing (r=0.16, p = 0.040) problems, and in particular with avoidant personality (r=0.22, p = 0.007) and ADHD (r=0.16, p =0.045) scales, although the latter did not survive the Bonferroni correction. The second adversity factor was not related to psychopathology. The correlations between single adversity measures and psychopathology can be found in Table S2.

### fMRI Results

#### Task Effect

A one-sample t-test was performed to identify brain regions involving EV and PE signaling. We found robust activation in key brain regions such as striatum (caudate, putamen, and nucleus accumbens) and medial prefrontal cortex during EV and PE signaling (Figure 3; p < 0.05, whole-brain FWE-corrected), which was compatible with a previous meta-analysis on neural correlates of reinforcement learning (7). Detailed list of brain regions showing activation/deactivation during EV and PE encoding can be found in the Supplementary Material S5 (Table S4-S7).

**Figure 3.**
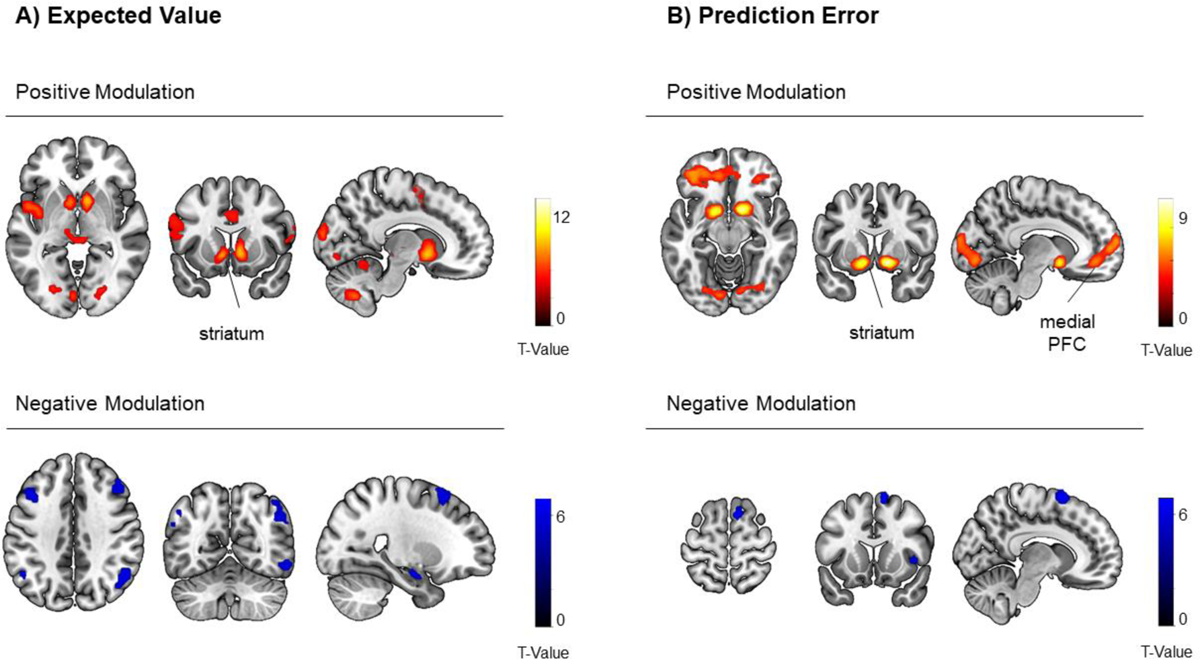
Expected value and prediction error signaling in the brain (p <0.05, whole-brain FWE corrected). Results were mapped on the brain surface using MRIcroGL toolbox (https://www.nitrc.org/projects/mricrogl).

### Adversity Effect

#### Multivariate Effects

The associations between adversity factors and EV signaling are shown in Figure 4. The first adversity factor, related to postnatal psychosocial adversities and prenatal smoking, was associated with lower EV activation in the vmPFC cluster extending to the left amygdala and left parahippocampal gyrus (p = 0.02, cluster level FWE-corrected; cluster size= 268, z= 3.38). However, the vmPFC cluster did neither survive the Bonferroni correction nor bootstrapping test, and should be considered preliminary.

**Figure 4.**
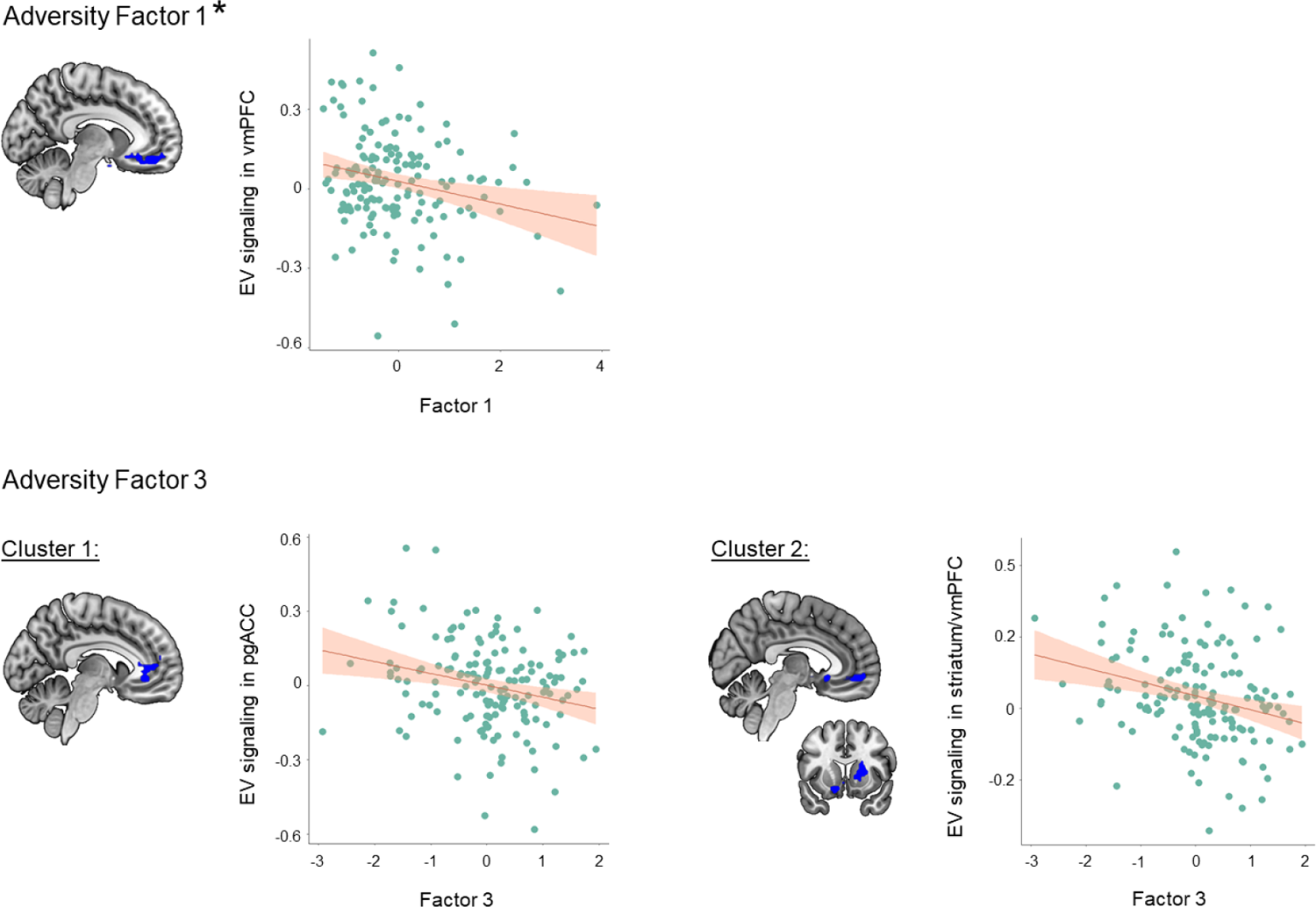
The associations between adversity factors and expected value representation. Blue color represents lower expected value signaling in the regions-of-interest. * The results did not survive after Bonferroni correction and bootstrapping test. Abbreviations: EV, expected value; pgACC, pregenual anterior cingulate cortex; vmPFC, ventromedial prefrontal cortex.

We did not find any significant cluster related to the second adversity factor.

The third adversity factor, reflecting mostly low maternal sensitivity, was associated with lower EV encoding in a large cluster including vmPFC, right striatum (caudate, putamen, nucleus accumbens), and right insula (p < 0.001, cluster level FWE-corrected; cluster size= 836, z= 4.24). A second cluster included pregenual ACC (p = 0.012, cluster level FWE-corrected; cluster size= 294, z= 3.97). These results remained significant after the Bonferroni correction and bootstrapping test. We did not find any significant PE signaling alteration for the adversity factors.

#### Specific Adversity Effects

To determine the specific contribution of each adversity measure to EV and PE encoding, we performed separate analyses in SPM. Two adversity measures were associated with significant alterations in EV and PE signaling (Figure 5). Lower maternal stimulation was associated with lower EV encoding in vmPFC as well as right striatum (caudate and putamen) and pregenual ACC (p < 0.001, cluster level FWE-corrected; cluster size= 1833, z= 4.61), thereby confirming the previous analysis based on adversity factors. Interestingly, obstetric adversity was related to lower PE representation in the pregenual ACC, vmPFC, and several prefrontal regions (p < 0.001, cluster level FWE-corrected; cluster size= 1342, z= 4.07). These results were robust after the Bonferroni correction and bootstrapping test. We did not find any significant alterations in EV and PE signaling for the other adversity measures.

**Figure 5.**
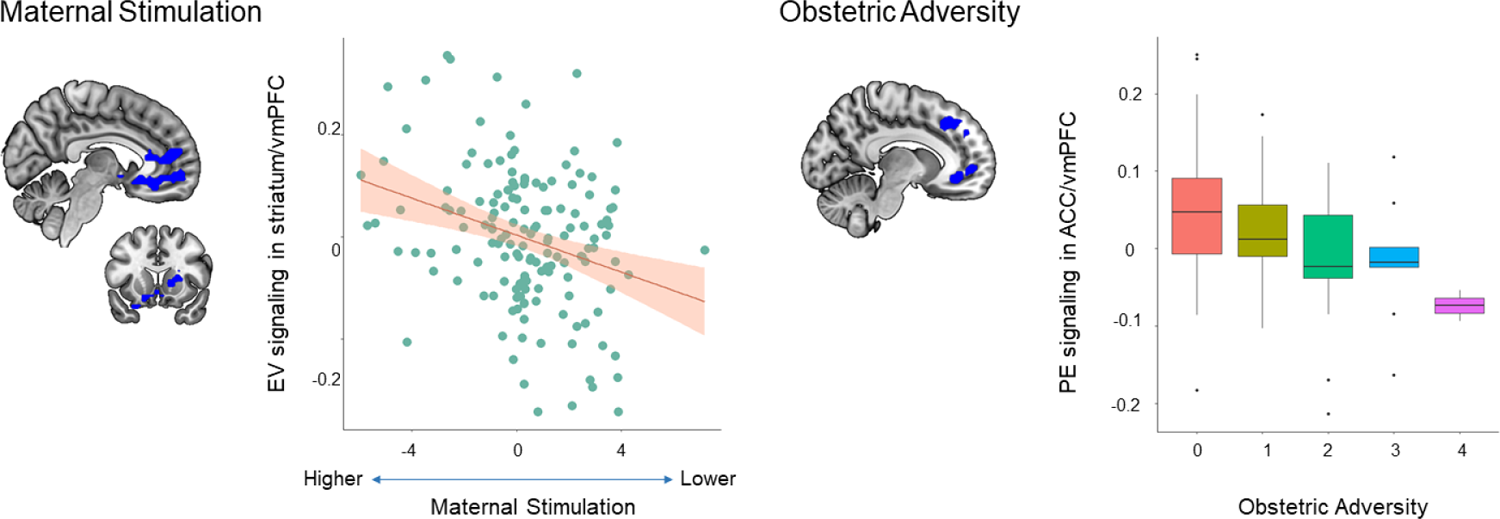
The associations between adversity measures and expected value/prediction error signaling. Maternal stimulation scores are reverse coded. Higher scores indicates lower maternal care. Blue color represents lower expected value and prediction error representation in the regions-of-interest. Abbreviations: EV, expected value; PE, prediction error; pgACC, pregenual anterior cingulate cortex; vmPFC, ventromedial prefrontal cortex.

Whole-brain results for adversity factors and single adversity measures are reported in the Supplementary Material (Table S8-S9).

#### Exploratory Analyses

As expected, family adversity measures highly correlated with each other, while the life events measures exhibited small-to-moderate correlations (Table S10-S11). Our exploratory analyses on the effects of psychosocial adversities occurring during different developmental periods (Supplementary Material S7) revealed that higher family adversity during infancy (T2) was linked to lower EV signaling in striatum, while family adversity during toddlerhood (T3) was related to lower EV encoding in the striatum and vmPFC (all p<0.01, cluster level FWE-corrected). Higher stressful life events during infancy (T2) and childhood (T4) and higher current stress (T11) was linked to lower EV signaling in vmPFC (all p <0.01, cluster level FWE-corrected), all of which did not survive Bonferroni correction. We did not identify any EV or PE abnormalities for stressful life events during adolescence and young adulthood.

#### Brain-Behavior Associations

The results for mediation analyses are shown in Figure 6. The first adversity factor predicted higher scores in all psychopathology measures including internalizing problems, externalizing problems, depression, anxiety, avoidant personality, somatic problems, ADHD, and antisocial personality subscales, and was related to lower EV representation in the vmPFC. Therefore, we run mediation analyses to see if EV representation in the vmPFC mediated the adversity-psychopathology association. Our results showed that the vmPFC partially mediated the relationship between the first adversity factor and ADHD symptoms (interaction effect (a*b)=0.21, CI= [0.04 0.50]).

**Figure 6.**
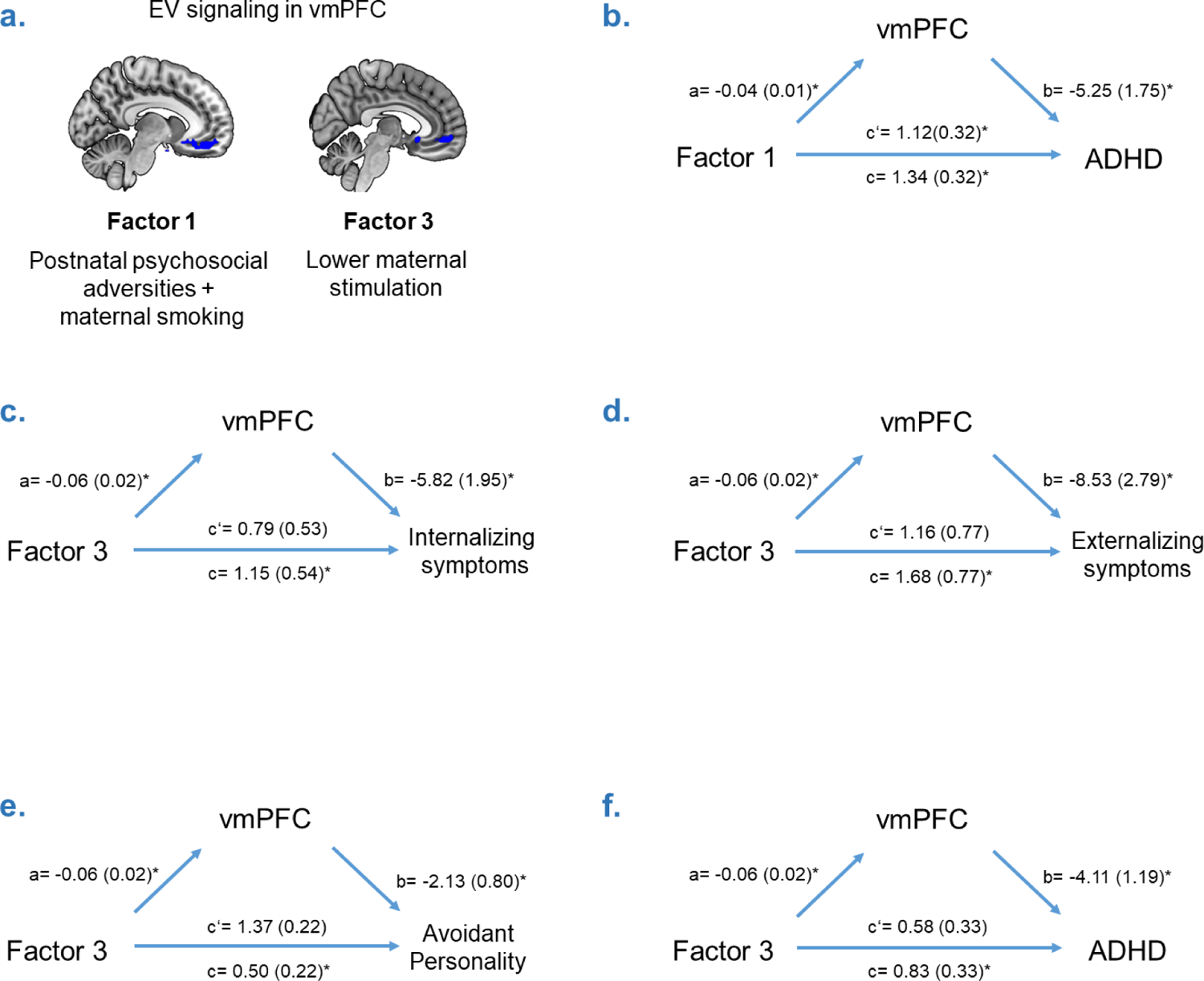
Mediation analysis. The association between adversity factors and expected value signaling in ventromedial prefrontal cortex (a). Mediation models (b-f). Each model included expected value signaling in ventromedial prefrontal cortex as a mediator (M) to explain the impact of adversity factors (X) on psychopathology symptoms (Y). Significant paths were shown with asterisk. Abbreviations: ADHD, attention deficit and hyperactivity disorder; vmPFC, ventromedial prefrontal cortex.

The third adversity factor predicted higher scores in internalizing problems, externalizing problems, avoidant personality, and ADHD subscales, and was associated with lower EV signaling in the pregenual ACC, vmPFC, and striatum. EV signaling in the vmPFC fully mediated the relationship between the third adversity factor and ASR internalizing (interaction effect= 0.35, CI= [0.09 0.75]), externalizing (interaction effect=0.51, CI= [0.15 1.06]), avoidant personality (interaction effect=0.13, CI= [0.02 0.30]) and ADHD (interaction effect=0.25, CI= [0.05 0.56]) scales.

Similar to these results, EV encoding in vmPFC was specifically associated with several psychopathology measures (See Supplementary Material S8 for correlation analyses). No association was found for other regions.

## Discussion

Capitalizing on data from a birth cohort, we investigated the specific and combined effect of lifespan adversities on EV and PE encoding. Our findings showed that adversities were associated with lower EV and PE signaling in the striatum and prefrontal cortex, and altered EV signaling further mediated the relationship between adversities and psychopathology. These results critically extend previous reports (21–23) by incorporating a developmental perspective, considering multiple prospectively assessed risk factors, and offering compelling evidence for enduring neurobiological and psychopathological consequences.

The findings regarding the three-factor adversity solution indicated that the third adversity factor reflecting lower maternal sensitivity was associated with a more extensive EV disruption in the brain, including vmPFC, ACC and striatum. This finding underscores the importance of caregiver-infant interaction on reward learning, which is essential for learning, exploration and normative brain development (5,6,49). Inconsistent caregiver behavior can create an unstable environment where the rewards are sparse and random, and thus impair the utilization of environmental information to optimize the behavior (5). On the other hand, consistent and good quality maternal care can buffer negative outcomes of adverse experiences on reward processing and learning. Indeed, a previous study from our lab showed that higher maternal stimulation was associated with increased striatum activation during reward anticipation in young adults with parental psychopathology (32). Therefore, implementing programs for promoting positive parenting in risk groups could help infants and children to develop normative behavioral and brain responses in reward processing and learning. Moreover, while the second adversity factor capturing common variance in obstetric adversity and prenatal maternal stress did not yield any EV or PE abnormality, specifically analyzing obstetric adversity showed disrupted PE signaling in ACC and vmPFC. Lastly, the first adversity factor informed by postnatal psychosocial adversities and prenatal smoking was associated with EV abnormalities in vmPFC. However, this result was not robust.

Moreover, our results indicated that the adversity measures collected very early in life (up to 3 months of age) can disrupt neural representation of EV and PE signaling. These results were also confirmed by our exploratory analyses on the timing effect of adversities (Supplementary Material S7), which revealed that family adversity experienced during infancy and toddlerhood had an impact on EV signaling in striatum and vmPFC. Several studies suggested that stressors occurring in early life are more likely to affect reward circuitry (5,17,50). Indeed, a previous study performing a sensitivity period analysis found that the striatum was sensitive to maltreatment that occurred between the ages of 0-4 years (51). We also identified EV abnormalities in individuals with higher current stress. However, this result was restricted to vmPFC. Taken together, these results may indicate that the striatum can be more sensitive to early life stress occurring in a caregiver context, as seen in lower maternal stimulation and adverse family environment in our study and maltreatment in a previous study (51), whereas stress over the lifespan may have an impact on the prefrontal cortex. Previous literature suggests that prefrontal cortex has a longer maturation window (52) and remarkable neural plasticity in adulthood (53), therefore, it might be susceptible to stress over the life course. However, more research is needed to make inferences about the sensitivity period for the reward network.

Importantly, we further showed that lower EV signaling in vmPFC was associated with psychopathology symptoms and mediated the relationship between adversity factors and psychopathology symptoms in internalizing and externalizing spectrum. These results are compatible with several previous studies that identified neural EV and PE abnormalities in depression (54), ADHD (3), substance abuse (55), generalized anxiety disorder (13), and conduct disorder (56). Taken together, these results indicate that disruptions in EV and PE signaling in the brain can increase the risk of developing psychopathology in individuals exposed to adversities via non-optimal decision-making processes.

Stress disrupts the dopaminergic system in animals (57) and humans (58,59), which is pivotal for EV and PE signaling. Indeed, there is a striking overlap between stress and reward neurobiology, which recruits the same brain regions including the ventral striatum, amygdala, and medial prefrontal cortex (5,49,50). Moreover, these regions are dense in terms of glucocorticoid receptors(60), which were found to be altered following early life stress exposure in animals (61) and humans (62). Therefore, it is not surprising that we identified that adversities were associated with disrupted EV and PE signaling in the reward network. Indeed, several previous studies reported functional (16–18,20,63,64), structural (65,66) and white matter tract (67,68) abnormalities in striatum and prefrontal cortex in individuals exposed to adversity. Disrupted EV and PE signaling in striatum and prefrontal cortex in individuals exposed to adversities may lead to impairments in several important skills such as subjective value representation, approach behavior, and risk/benefit assessment (16,18). These skills might especially be important to adapt rapidly changing environments that require flexibility to face new challenges as we have nowadays experienced with the COVID-19 pandemic, wars and climate change.

### Limitations

Here, we investigated the combined and specific effects of developmental adversities on neural correlates of reinforcement learning using a longitudinal design with several prospective adversity measures in a relatively large well-phenotyped sample. However, the current study has also some limitations. First, we did not investigate the differential neural responses to reward and punishment prediction errors to increase power in the statistical analysis. However, several studies suggest that reward and loss networks are similar (69,70).

Second, our sample contained mostly healthy participants. Although our results indicated vulnerability direction given the relationship with psychopathology, the limited variation in psychopathology symptoms warrants further validation in clinical samples.

## Conclusions

In conclusion, we showed that adversities across the development were linked to altered neural EV signaling in the reward network in adulthood, and neural alterations mediated the relationship between adversities and internalizing and externalizing psychopathology. These results indicate that developmental risk factors are related to impaired neural processing of reward-related cues and contingencies, which in turn, are linked to enhancing the risk of developing psychopathology. Therefore, it is very important to develop preventative strategies for individuals at risk to buffer negative outcomes of adverse experiences on the brain and behavior.

## Acknowledgements

The authors would like to thank the participants for their continued participation in the Mannheim Study of Children at Risk and the reviewers for their constructive comments and suggestions. NEH gratefully acknowledges grant support from the German Research Foundation (grant numbers DFG HO 5674/2-1, GRK2350/1) and in the framework of the Radboud Excellence Fellowship. TB gratefully acknowledges grant support by the German Federal Ministry of Education and Research (01EE1408E ESCAlife; FKZ 01GL1741[X] ADOPT; 01EE1406C Verbund AERIAL; 01EE1409C Verbund ASD-Net; 01GL1747C STAR; 01GL1745B IMAC-Mind), by the German Research Foundation (TRR 265/1), by the Innovative Medicines Initiative Joint Undertaking (IMI JU FP7 115300 EU-AIMS; grant 777394 EU-AIMS-2-TRIALS) and the European Union – H2020 (Eat2beNICE, grant 728018; PRIME, grant 847879).TUH is supported by a Sir Henry Dale Fellowship (211155/Z/18/Z; 211155/Z/18/B; 224051/Z/21) from Wellcome & Royal Society, the Medical Research Foundation, and a Philip Leverhulme Prize from the Leverhulme Trust (PLP-2021-040). TUH is also supported by the Carl-Zeiss-Stiftung. The Max Planck UCL Centre is a joint initiative supported by UCL and the Max Planck Society. The Wellcome Centre for Human Neuroimaging is supported by core funding from the Wellcome Trust (203147/Z/16/Z).

