## Supplementary Material for "Early life adversities affect expected value signaling in the adult brain"

† These authors contributed equally.

### S1. Exclusion Criteria for fMRI

Two hundred and fifty six participants (67%) agreed to participate in the study at the last assessment wave. Functional magnetic resonance imaging (fMRI) data for the passive avoidance task was available for 170 participants. Fourteen participants were excluded due to inefficient coverage of the brain surface (n=3), excessive head motion (n=4; > 3 mm in transition or 3 degrees in rotation), no understanding of the task (n=1) and technical problems during the task administration (n=6). Included participants did not differ in terms of sex and adversities except the prenatal maternal smoking. Excluded participants had higher prenatal maternal smoking scores compared to the included participants ($t$ (168)=2.82, $p < 0.01$).

### S2. Adversity Measures across Development

**Prenatal Maternal Smoking**

We measured maternal smoking during pregnancy by a standardized interview conducted with mothers at the 3-month assessment (1). Mothers were asked about their daily cigarette consumption (1= no, 2= up to 5 per day, 3= more than 5 per day). The score range is 1-5. Of the mothers of participants, 116 (74.36%) were nonsmokers, 17 (10.9%) reported smoking 1 to 5 cigarettes per day, and 23 (14.74%) reported smoking more than five cigarettes per day.

**Prenatal Maternal Stress**

Prenatal maternal stress was measured at the first assessment wave (age of 3 months) via a standardized parent interview conducted by trained interviews (2). Mothers answered 11 questions covering negative (e.g., ‘Did you have mood swings/ a depressed mode’) and negatively coded positive experiences (e.g., ‘Did you look forward to having a baby’) during pregnancy. Additionally, mothers were asked about the timing of these experiences (the first and the second/third trimesters). We here used the prenatal maternal stress scores for the second/third trimesters since the effect of mid- and late pregnancy on offspring behavior was found to be the largest (3). The items were coded dichotomously based on the presence of experience. The score range is 0-8. Higher scores indicated higher prenatal maternal stress.

**Obstetric Adversity**

Obstetric adversity scores were calculated according to the degree of obstetric complications based on medical reports (4). The score range is 0-4 (0= no risk, 1-2=moderate risk, 3-4= high risk). Sixty-two infants (39.75%) were born full-term, had normal birth weights and had no medical complications. Ninety-four infants had moderate (53.2%; e.g., preterm birth, preterm labor) to high (7.05%; e.g., low birth weight (< 1500 kg), neonatal complications) biological risk.

**Maternal Stimulation**

Videotapes of a 10-min standardized nursing and playing situation between mothers and their 3-month-olds were recorded and evaluated by trained raters (κ>0.83) using a modified version of the categorical system for micro-analysis of the early mother-child interaction (5). Raters were blind to parental and child risk status. We coded the presence or absence of nine measures of mother-infant interaction behavior in 120 five-second intervals. Maternal stimulation included all attempts (vocal, facial or motor) to attract the infant´s attention or to establish contact with him/her. It was coded as present when the baby was gazing at the mother or the behaviors were directed to the child. The scores were z-transformed, and recoded such that higher scores indicated lower maternal stimulation.

**Family Adversity**

Family adversity scores (Laucht et al., 2000) were calculated based on the presence of 11 adverse family factors covering characteristics of the parents (e.g., low educational level, psychiatric history), the partnership (e.g., marital discord), and the family environment (e.g., overcrowding) in the period from T1 (age of 3 months) to T5 (age of 11 years). The score range is 0-10 for the current sample. Higher scores indicated higher psychosocial risk.

**Childhood Trauma Questionnaire**

Childhood Trauma Questionnaire (CTQ) is a self-reported measure of traumatic experiences that occurred in childhood (6). We here used the German version of CTQ (7) which includes 28 items covering five subscales: emotional abuse, emotional neglect, physical abuse, physical neglect, and sexual abuse.  Responses are quantified on a 5-point Likert scale (1= ‘never true’, 5= ‘very often true’). The total score range is 25-87. We here used total CTQ scores (M=30.84, SD=8.58) as a measurement of adversity.

**Stressful Life Events**

We measured stressful life events from the first assessment wave (age of 3 months) until the last assessment wave (age of 33 years) using a modified version of the Munich Event List [MEL] (8). The MEL is an interview procedure for assessing acute and chronic as well as positive and negative stressors. It covers items regarding parental socio-economic disadvantages, negative health outcomes, and living and environmental conditions. Parents rated the MEL up to the seventh assessment wave (age of 15 years). Starting at the 15-year assessment, participants rated stressful life events themselves. Here, we calculated a sum score based on separate Z-transformed scores from T1 (age of 3 months) to time T11 (age of 33 years). Higher scores indicated higher stressful life events.

**Table S1.** Correlations between adversity measures.

| | Maternal Stress | Maternal Smoking | Maternal Stimulation | Obstetric Adversity | Family Adversity | Childhood Trauma Questionnaire | Stressful Life Events |
|---|---|---|---|---|---|---|---|
| Maternal Stress | - | 0.09 [b] | -0.04 [b] | **0.20*[b]** | **0.26** [b]** | 0.12 [b] | 0.15 [b] |
| Maternal Smoking | | - | -0.01 [b] | -0.02 [b] | **0.32*** [b]** | 0.15 [b] | **0.39*** [b}** |
| Maternal Stimulation | | | - | -0.02 [b] | **-0.24**[a]** | -0.14 [b] | **-0.18*[a]** |
| Obstetric Adversity | | | | - | -0.09 [b] | -0.02 [b] | -0.01 [b] |
| Family Adversity | | | | | - | **0.33*** [b]** | **0.56*** [a]** |
| Childhood Trauma Questionnaire | | | | | | - | **0.41*** [b]** |
| Stressful Life Events | | | | | | | - |

* $p < 0.05$  ** $p < 0.01$  *** $p < 0.001$. a= Pearson's correlation test, b= Spearman's correlation test. Significant correlations were shown in bold font.

**Table S2.** Spearman's correlations between adversity and psychopathology measures.

| | Internalizing Symptoms | Externalizing Symptoms | Depression | Anxiety | Avoidant Personality | Somatic Problems | ADHD | Antisocial Personality |
|---|---|---|---|---|---|---|---|---|
| Maternal Stress | 0.18 * | - | - | - | - | - | - | - |
| Maternal Smoking | - | 0.19 * | 0.22** | - | - | - | - | - |
| Maternal Stimulation | - | - | - | - | -0.16* | - | - | - |
| Obstetric Adversity | - | - | - | - | - | - | - | - |
| Family Adversity | 0.20* | **0.22*** | **0.26**** | 0.18* | - | - | 0.20* | 0.19* |
| Childhood Trauma Questionnaire | **0.38***** | **0.48***** | **0.44***** | **0.39***** | **0.37***** | 0.16* | **0.31***** | **0.38***** |
| Stressful Life Events | **0.36***** | **0.36***** | **0.39***** | **0.33***** | 0.21** | 0.21** | **0.28***** | **0.34***** |

* p < 0.05 ** p < 0.01 *** p < 0.001. The correlations survived after the Bonferroni correction were shown in bold font. Abbreviations: ADHD, Attention deficit and hyperactivity disorder.

### S3. Preprocessing Pipeline

The first five scans were discarded to allow for equilibration of the magnetic field. The preprocessing steps included slice timing correction of volumes to the middle slice, realignment to the first volume using a rigid body linear transformation, structural and functional image co-registration, segmentation, normalization to the Montreal Neurological Institute (MNI) template, and smoothing using a kernel with a full-width half-maximum of 8 mm.

#### S4. Computational Modelling

To understand the observed behavior of participants during decision-making, we compared the variations of five Rescorla-Wagner models (See Table S3). The first model included three parameters, V0 (i.e., expected value for the first trial), $\alpha$ (learning rate) and a standard SoftMax function parameter $\beta$ (i.e., stochasticity/inverse temperature). The second and third models included an extended SoftMax function parameter, $\pi$ (i.e., constant pressing bias) and $\pi$_t (i.e., pressing bias decreasing with time) respectively, in addition to the above-mentioned parameters. The fourth model was similar to the third model but included separate learning rates for positive and negative prediction errors instead of one common learning rate (9). The fifth model had two pressing bias ($\pi$_t) parameters for each block instead of one pressing bias parameter across the task and varied in terms of learning rate (common vs. separate). Each model was fitted to each participant's data, and the best model was determined using Akaike and Bayesian Information Criteria (Table S3, Figure S1 and S2). The winning model was the second with four free parameters (V0, $\alpha$, $\beta$, and $\pi$_t; Figure S3).

**Table S3.** Model descriptions.

| Model | NTrials | nLL | NParams | BIC | AIC |
|---|---|---|---|---|---|
| Model0_alpha | 112 | 57.47 | 1 | 119.65 | 116.93 |
| Model0_alpha_beta | 112 | 53.94 | 2 | 117.33 | 111.89 |
| Model0_beta | 112 | 55.67 | 1 | 116.05 | 113.33 |
| Model0_v0 | 112 | 57.49 | 1 | 119.69 | 116.97 |
| Model0_v0_alpha | 112 | 52.87 | 2 | 115.17 | 109.74 |
| Model0_v0_alpha_beta | 112 | 48.79 | 3 | 111.71 | 103.56 |
| Model0_v0_beta | 112 | 51.03 | 2 | 111.49 | 106.05 |
| Model1_alpha_beta_pi | 112 | 48.48 | 3 | 111.11 | 102.95 |
| Model1_alpha_pi | 112 | 51.28 | 2 | 111.99 | 106.55 |
| Model1_v0_alpha_beta_pi | 112 | 45.83 | 4 | 110.54 | 99.66 |
| Model1_v0_alpha_pi | 112 | 49.03 | 3 | 112.21 | 104.05 |
| Model2_alpha_beta_pi_t | 112 | 48.22 | 3 | 110.60 | 102.44 |
| Model2_alpha_pi_t | 112 | 51.20 | 2 | 111.84 | 106.40 |
| **Model2_v0_alpha_beta_pi_t** | **112** | **43.11** | **4** | **105.10** | **94.23** |
| Model2_v0_alpha_pi_t | 112 | 47.98 | 3 | 110.13 | 101.97 |
| Model3_2alpha | 112 | 52.07 | 2 | 113.58 | 108.14 |
| Model3_2alpha_beta | 112 | 49.35 | 3 | 112.85 | 104.70 |
| Model3_2alpha_beta_pi_t | 112 | 46.46 | 4 | 111.80 | 100.92 |
| Model3_2alpha_pi_t | 112 | 48.67 | 3 | 111.50 | 103.34 |
| Model3_v0_2alpha_beta | 112 | 46.88 | 4 | 112.64 | 101.76 |
| Model3_v0_2alpha_beta_pi_t | 112 | 43.67 | 5 | 110.52 | 96.93 |
| Model3_v0_2alpha_pi_t | 112 | 45.80 | 4 | 110.49 | 99.61 |
| Model4_2alpha_beta_2pi_t | 112 | 43.71 | 5 | 111.02 | 97.43 |
| Model4_alpha_beta_2pi_t | 112 | 45.82 | 4 | 110.52 | 99.65 |
| Model4_v0_2alpha_beta_2pi_t | 112 | 41.51 | 6 | 111.33 | 95.02 |
| Model4_v0_alpha_beta_2pi_t | 112 | 43.71 | 5 | 111.00 | 97.41 |

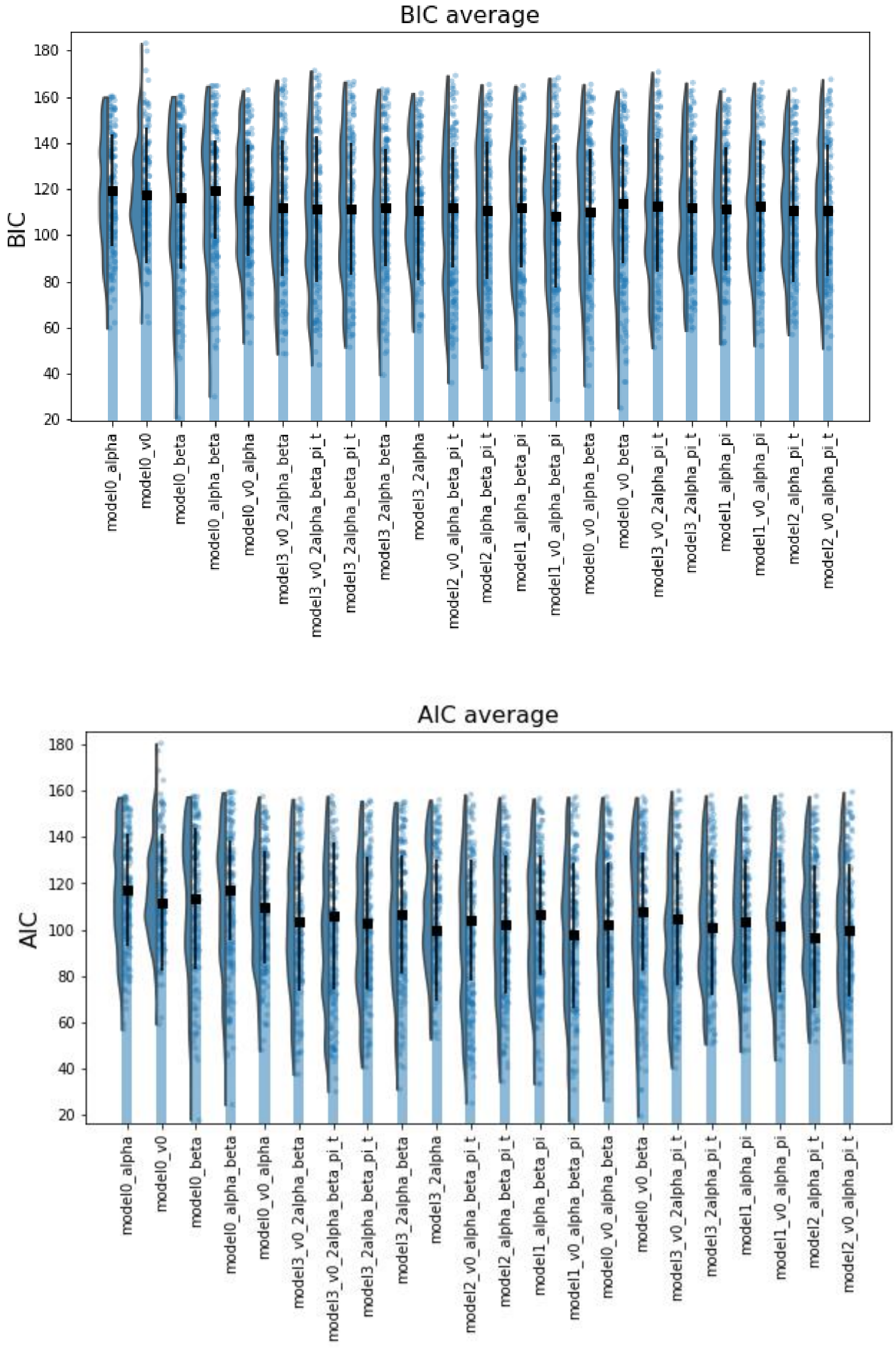

**Figure S1.** Model Comparison based on Bayesian and Akaike Information Criteria.

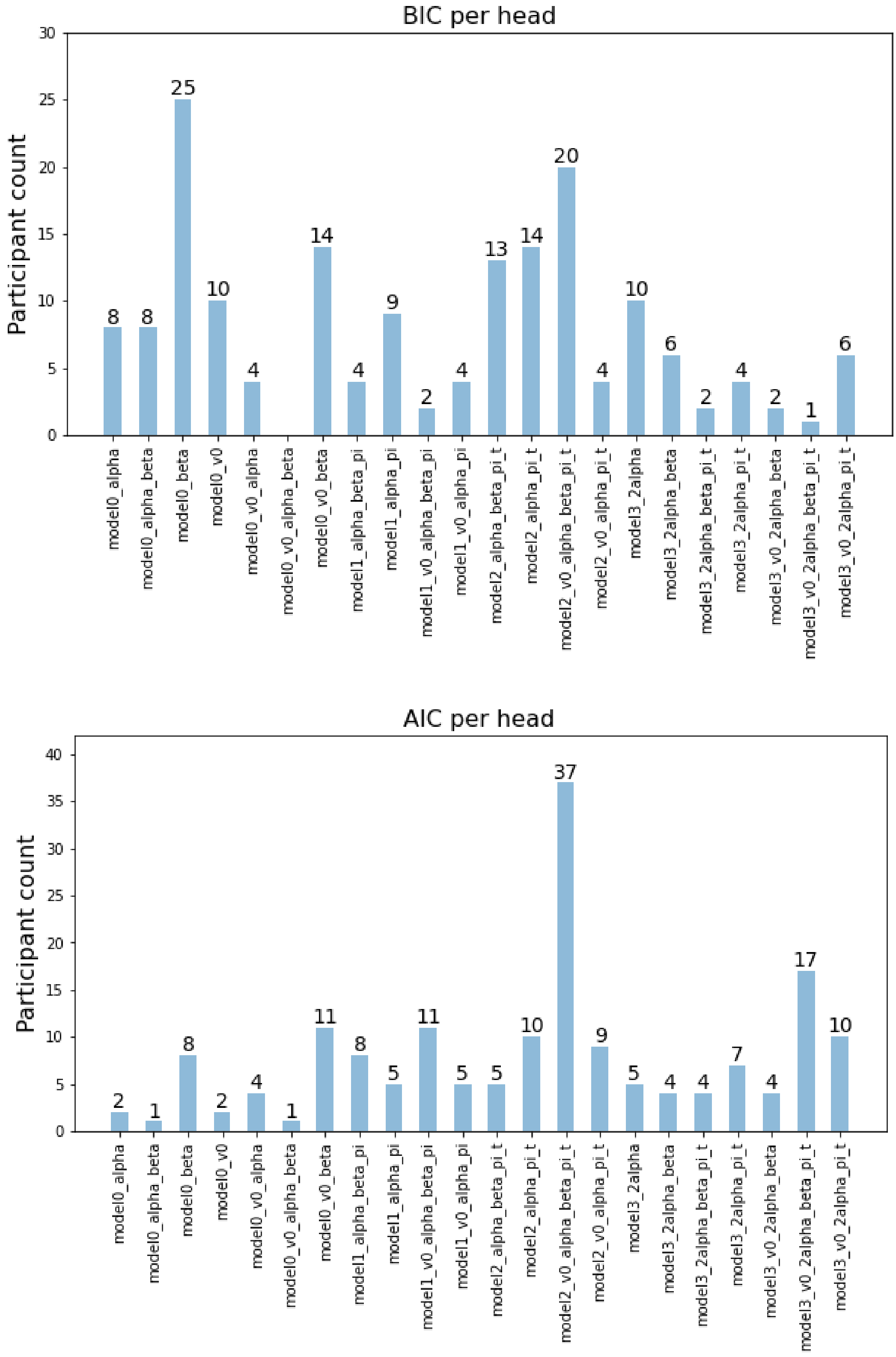

**Figure S2.** Model Comparison per head. The figure shows the winning models per participant.

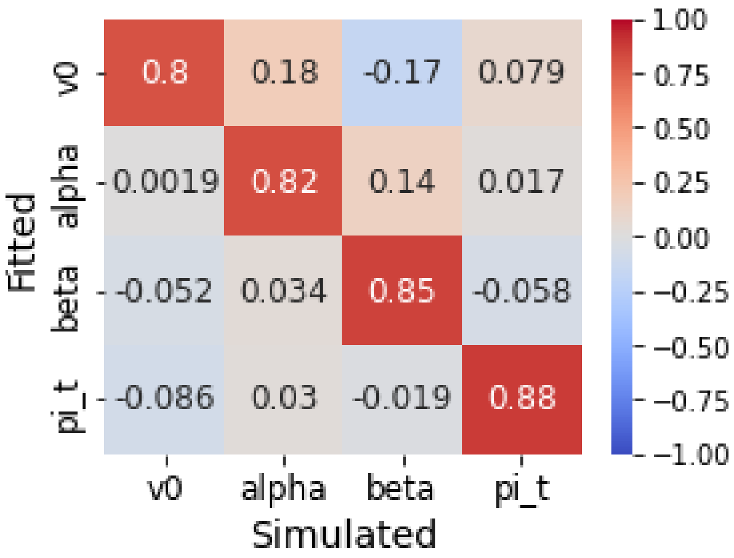

**Figure S3.** Parameter recovery matrix for the winning model. The matrix shows the correlations between simulated and fitted parameters. The diagonal entries of the matrix are near to one, whereas the most of the off-diagonal entries are close to zero, indicating that parameters of the model were identifiable.

**Expected Value and Prediction Error Calculation**

On each trial t, we calculated PE using the following formula.

$$PE_{(t)} = FB_{(t)} - EV_{(t)}$$

In this formula, the PE for the current trial is equaled to the feedback value minus the EV for the current trial. For the first trial, EV was set to subject-specific parameter estimation obtained from the winning model and was then updated with the following formula.

$$EV_{(t)}=EV_{(t-1)} + (\alpha * PE_{(t-1)})$$

In this formula, the EV for the current trial is equaled to the EV for the previous trial plus the PE for the previous trial multiplied by the learning rate. The learning rate was set to the subject-specific parameter estimation obtained from the winning model.

We then calculate the probability of hitting ($P_{hit}$) in the SoftMax function using the following formula:

$$V_{hit} = EV + \pi\_t * shrink$$

$$P_{hit} = 1 / (1 + \exp(\beta *(-V_{hit})))$$

Shrink is a parameter that starts high at the beginning of each block and then decreases across the trials, which is defined as the total number of trials minus the current trial number divided by the total number of trials ((56 – current trial number)/56). It captures the fact that participants are more keen to hit at the beginning of a block.

#### S5. Neural Correlates of Expected Value and Prediction Error

We identified robust activation in key brain regions during expected value (EV) and prediction error (PE) signaling such as striatum (caudate, putamen, and nucleus accumbens) and medial prefrontal cortex ($p < 0.05$, whole-brain FWE-corrected). During the cue phase, we found higher EV signaling in the bilateral striatum (caudate, putamen, and nucleus accumbens), midbrain, pre- and postcentral gyrus, supplementary motor area, insula, occipital cortex, and cerebellum (Table S4, Table S5). We also found lower EV encoding in bilateral middle frontal gyrus, angular gyrus, left occipital pole, right amygdala/hippocampus, and right inferior temporal cortex (Table S4). During the feedback phase, we found higher PE representation in the striatum (bilateral putamen, bilateral nucleus accumbens, and right caudate), orbitofrontal cortex, superior, medial, and inferior frontal gyrus, occipital cortex, left inferior parietal cortex, and cerebellum (Table S6, Table S7) and  lower PE representation in the right supplementary motor area and right insula during the feedback phase (Table S6).

**Table S4.** Peak coordinates of expected value signaling across the whole-brain.

| Regions | Hemisphere | Cluster Size | T | MNI Coordinates [x y z] | | |
|---|---|---|---|---|---|---|
| **Positive Modulations** | | | | | | |
| Postcentral gyrus | L | 3123 | 13.99 | -54 | -22 | 47 |
| Precentral gyrus | | | | | | |
| Supplementary motor area | | | | | | |
| Superior occipital gyrus | R | 227 | 10.34 | 27 | -91 | 20 |
| Cerebellum | R | 97 | 9.85 | 15 | -64 | -46 |
| Striatum | R | 121 | 9.79 | 9 | 8 | -4 |
| Cerebellum | R | 175 | 9.52 | 21 | -55 | -22 |
| Lingual gyrus | L | 188 | 9.38 | -9 | -85 | -7 |
| Striatum | L | 104 | 8.75 | -9 | 8 | -7 |
| Insula | R | 144 | 8.63 | 42 | -1 | 11 |
| Thalamus | L | 120 | 8.09 | -15 | -19 | 8 |
| Postcentral gyrus | R | 123 | 7.88 | 54 | -19 | 20 |
| Lingual gyrus | R | 94 | 7.70 | 24 | -79 | -7 |

**Table S4** (continued)

| | | | | | | |
|---|---|---|---|---|---|---|
| Superior occipital gyrus | L | 104 | 7.47 | -18 | -94 | 17 |
| Insula | L | 21 | 6.59 | -30 | 23 | 8 |
| Thalamus | R | 10 | 5.57 | 6 | -16 | 2 |
| **Negative Modulations** | | | | | | |
| Angular gyrus | R | 148 | 6.62 | 54 | -55 | 32 |
| Occipital pole | L | 39 | 6.51 | -24 | -100 | -7 |
| Middle frontal gyrus | R | 271 | 6.50 | 42 | 29 | 44 |
| Middle frontal gyrus | L | 229 | 6.34 | -39 | 14 | 53 |
| Right amygdala | R | 22 | 6.15 | 27 | -7 | -19 |
| Inferior temporal gyrus | R | 37 | 5.98 | 54 | -61 | 41 |
| Angular gyrus | L | 45 | 5.77 | -48 | -61 | 41 |

$p < 0.05$ (whole-brain FWE corrected, cluster size >= 10)

**Table S5.** Peak coordinates of expected value signaling in the regions of interest.

| Regions | Hemisphere | Cluster Size | T | MNI Coordinates [x y z] | | |
|---|---|---|---|---|---|---|
| **Positive Modulations** | | | | | | |
| Caudate | L | 12 | 7.25 | -6 | 8 | -1 |
| | R | 31 | 9.22 | 9 | 11 | -1 |
| Putamen | L | 14 | 7.61 | -15 | 5 | -10 |
| Nucleus accumbens | L | 30 | 8.75 | -9 | 8 | -7 |
| | R | 34 | 9.79 | 9 | 8 | -4 |
| Anterior cingulate cortex | L+R | 6 | 5.62 | 0 | 8 | 29 |

$p < 0.05$ (whole-brain FWE corrected)

**Table S6.** Peak coordinates of prediction error signaling across the whole brain.

| Regions | Hemisphere | Cluster Size | T | MNI Coordinates [x y z] | | |
|---|---|---|---|---|---|---|
| **Positive Modulations** | | | | | | |
| Striatum (caudate, nucleus accumbens, putamen) | R | 112 | 11.78 | 12 | 8 | -10 |
| Striatum (caudate, nucleus accumbens, putamen) | L | 99 | 10.17 | -12 | 5 | -10 |
| Cerebellum | R | 364 | 9.63 | 39 | -64 | -40 |
| Middle occipital gyrus | R | 1155 | 9.10 | 21 | -94 | 11 |
| Middle frontal gyrus | L | 896 | 7.92 | -36 | 44 | -10 |
| Orbitofrontal cortex | | | | | | |
| Superior frontal gyrus | L | 141 | 7.18 | -21 | 32 | 47 |
| Inferior parietal lobe | L | 251 | 6.87 | -48 | -46 | 47 |
| Cerebellum | L | 143 | 6.78 | -39 | -70 | -34 |
| Inferior orbital gyrus | R | 37 | 6.41 | 30 | 41 | -10 |
| Striatum (putamen) | L | 64 | 6.35 | -30 | -13 | 8 |
| Posterior cingulate cortex | L | 31 | 5.98 | -3 | -31 | 35 |

**Table S6** (continued)

| | | | | | | |
|---|---|---|---|---|---|---|
| Striatum (caudate) | R | 13 | 5.71 | 15 | 11 | 20 |
| Precentral gyrus | R | 34 | 5.62 | 3 | -28 | 62 |
| Striatum (putamen) | R | 14 | 5.49 | 33 | -4 | 2 |
| **Negative Modulations** | | | | | | |
| Superior frontal gyrus | R | 53 | 6.97 | 12 | 11 | 65 |
| Insula | R | 16 | 5.66 | 45 | 11 | 2 |

$p < 0.05$ (whole-brain FWE corrected, cluster size >= 10)

**Table S7.** Peak coordinates of prediction signaling in the regions of interest.

| Regions | Hemisphere | Cluster Size | T | MNI Coordinates [x y z] | | |
|---|---|---|---|---|---|---|
| **Positive Modulations** | | | | | | |
| Putamen | L | 68 | 10.04 | -15 | 5 | -10 |
| | R | 40 | 10.19 | 18 | 8 | -10 |
| Nucleus accumbens | L | 17 | 10.17 | -12 | 5 | -10 |
| | R | 32 | 11.78 | 12 | 8 | -10 |
| Anterior cingulate cortex | L | 33 | 6.18 | -6 | 53 | -1 |
| Ventromedial prefrontal cortex | L+R | 115 | 6.65 | -9 | 44 | -10 |

$p < 0.05$ (whole-brain FWE corrected)

#### S6. Effect of Adverse Experiences on Expected Value and Prediction Error Signaling at Whole-Brain

**Table S8.** The associations between the adversity factors, and expected value and prediction error signaling ($p < 0.05$, cluster level FWE corrected).

| | Contrast | Cluster | k | T | MNI Coordinates | | |
|---|---|---|---|---|---|---|---|
| Factor 1 | - | - | | | | | |
| Factor 2 | PE negative | Lingual gyrus R + L | 2012 | 4.57 | 24 | -67 | 17 |
| | | Calcarine R + L | | | | | |
| | | Middle temporal R | | | | | |
| | | Precuneus R + L | | | | | |
| | | Fusiform gyrus R + L | | | | | |
| Factor 3 | EV negative | Superior temporal gyrus L | 1613 | 4.34 | -54 | -1 | -7 |
| | | Postcentral gyrus L | | | | | |
| | | Precentral gyrus L | | | | | |
| | | Supramarginal gyrus L | | | | | |
| | | Inferior Parietal lobe L | | | | | |
| | | Insula L | | | | | |
| | | Thalamus L | | | | | |
| | | Putamen L | | | | | |

Abbreviations: EV, expected value; k, cluster size; PE, prediction error.

**Table S9.** The associations between adversity measures, and expected value and prediction error signaling (p <0.05, cluster level FWE corrected).

| | Contrast | Cluster | k | T | MNI Coordinates | | |
|---|---|---|---|---|---|---|---|
| CTQ | EV negative | Postcentral gyrus L | 419 | 3.80 | -48 | -28 | 41 |
| | | Supramarginal gyrus L | | | | | |
| | | Inferior parietal lobe L | | | | | |
| Maternal Stress | PE negative | Calcarine R | 381 | 4.28 | 27 | -67 | 14 |
| | | Lingual gyrus R | | | | | |
| | | Precuneus R | | | | | |
| | | Calcarine L | 282 | 3.32 | -18 | -61 | 8 |
| | | Lingual L | | | | | |
| | | Cuneus L | | | | | |
| Maternal Stimulation* | EV positive | Postcentral gyrus L | 518 | 4.49 | -15 | -10 | 80 |
| | | Superior parietal lobe L | | | | | |
| | | Superior frontal lobe L | | | | | |
| | EV positive | Superior temporal lobe L | 384 | 4.30 | -54 | -1 | -7 |
| | | Postcentral gyrus L | | | | | |
| | | Supramarginal gyrus L | | | | | |
| | EV positive | Middle temporal lobe L | 614 | 4.16 | -45 | -55 | 17 |
| | | Middle occipital lobe L | | | | | |
| | | Superior temporal lobe L | | | | | |
| | | Angular gyrus L | | | | | |
| | | Putamen L | | | | | |

**Table S9** (continued)

| | | | | | | | |
|---|---|---|---|---|---|---|---|
| | EV positive | Postcentral gyrus R | 250 | 3.87 | 54 | -25 | 56 |
| | | Middle frontal gyrus R | | | | | |
| | | Superior frontal gyrus R | | | | | |
| Obstetric Adversity | PE negative | Cuneus R | 1608 | 4.16 | 30 | -76 | 11 |
| | | Calcarine R | | | | | |
| | | Lingual gyrus R | | | | | |
| | | Middle temporal lobe R | | | | | |
| | | Fusiform R | | | | | |
| | | Cerebellum R | | | | | |
| | PE negative | Middle occipital gyrus L | 1019 | 4.11 | -27 | -52 | -13 |
| | | Middle temporal lobe L | | | | | |
| | | Lingual gyrus L | | | | | |
| | | Fusiform gyrus L | | | | | |
| | PE negative | Postcentral gyrus R | 316 | 3.97 | 3 | -25 | 65 |
| | | Supplementary motor area R | | | | | |
| | | Precuneus L | | | | | |

Abbreviations: EV, expected value; k, cluster size; PE, prediction error. * Higher maternal stimulation scores indicated higher maternal care.

#### S7. Exploratory Analyses: Timing Effect of Adversities

To investigate the timing effect of prospectively collected adversities on EV and PE signaling, we conducted a series of multiple regression analyses. In total, we performed five tests for family adversity (T1-T5) and eleven tests for stressful life events (T1-T11). Stressful life events measured at the last assessment wave (T11) was used as a measure of current stress.

Family adversity measures were not normally distributed and showed moderate to high correlations with each other (Table S10). Stressful life events were roughly normally distributed and showed no to moderate correlations with each other (Table S11). None of the correlations between variables caused to multicollinearity problem ($r > 0.8$).

**Table S10.** The Spearman's correlations between family adversity measures.

| | T1 | T2 | T3 | T4 | T5 |
|---|---|---|---|---|---|
| T1 | - | .68*** | .61*** | .52*** | .49*** |
| T2 | | - | .68*** | .53*** | .46*** |
| T3 | | | - | .66*** | .56*** |
| T4 | | | | - | .68*** |
| T5 | | | | | - |

*** $p < 0.001$

**Table S11.** The Pearson's correlations between stressful life events measures.

| | T1 | T2 | T3 | T4 | T5 | T6 | T7 | T8 | T9 | T10 | T11 |
|---|---|---|---|---|---|---|---|---|---|---|---|
| T1 | - | .49*** | .37*** | .20* | .31*** | .24** | .19* | .32*** | .11 | .12 | .05 |
| T2 | | - | .54*** | .32*** | .27** | .24** | .22** | .30*** | .27** | .15 | .21** |
| T3 | | | - | .37*** | .22** | .23** | .13 | .18* | .24** | .18* | .11 |
| T4 | | | | - | .38*** | .24** | .09 | .14 | .20* | .09 | .09 |
| T5 | | | | | - | .46*** | .40*** | .18* | .23** | .04 | .07 |
| T6 | | | | | | - | .26** | .10 | .13 | .08 | -0.01 |
| T7 | | | | | | | - | .38*** | .36*** | .18* | .24** |
| T8 | | | | | | | | - | .43*** | .35*** | .39*** |
| T9 | | | | | | | | | - | .37*** | .28*** |
| T10 | | | | | | | | | | - | .45*** |
| T11 | | | | | | | | | | | - |

*p <0.05 **p<0.01 ***p<0.001

Our exploratory analyses (Figure S4) revealed that higher family adversity at the age of 2 years was linked to lower EV encoding in the bilateral striatum (p=0.003, cluster level FWE-corrected, k=370, z=4.20). Moreover, lower EV signaling in vmPFC extending to the striatum was associated with higher family adversity at the age of 4 years (p=0.004, cluster level FWE-corrected, k=361, z=4.12). These results survived after the Bonferroni correction (p < 0.05/5=0.01). Higher stressful life events between the ages of 3 months and 2 years (p=0.005, cluster level FWE-corrected, k=350, z=3.58) and between the ages of 4 years and 8 years (p=0.037, cluster level FWE-corrected, k=238, z=3.78) were related to lower EV encoding in vmPFC. Current life stress was also associated with lower EV representation in vmPFC (p=0.009, cluster level FWE-corrected, k=311, z=4.58). However, these results did not survive after the Bonferoni correction (p > 0.05/11=0.004). We did not identify any significant cluster during EV and PE signaling for life events covering adolescence and young adulthood periods.

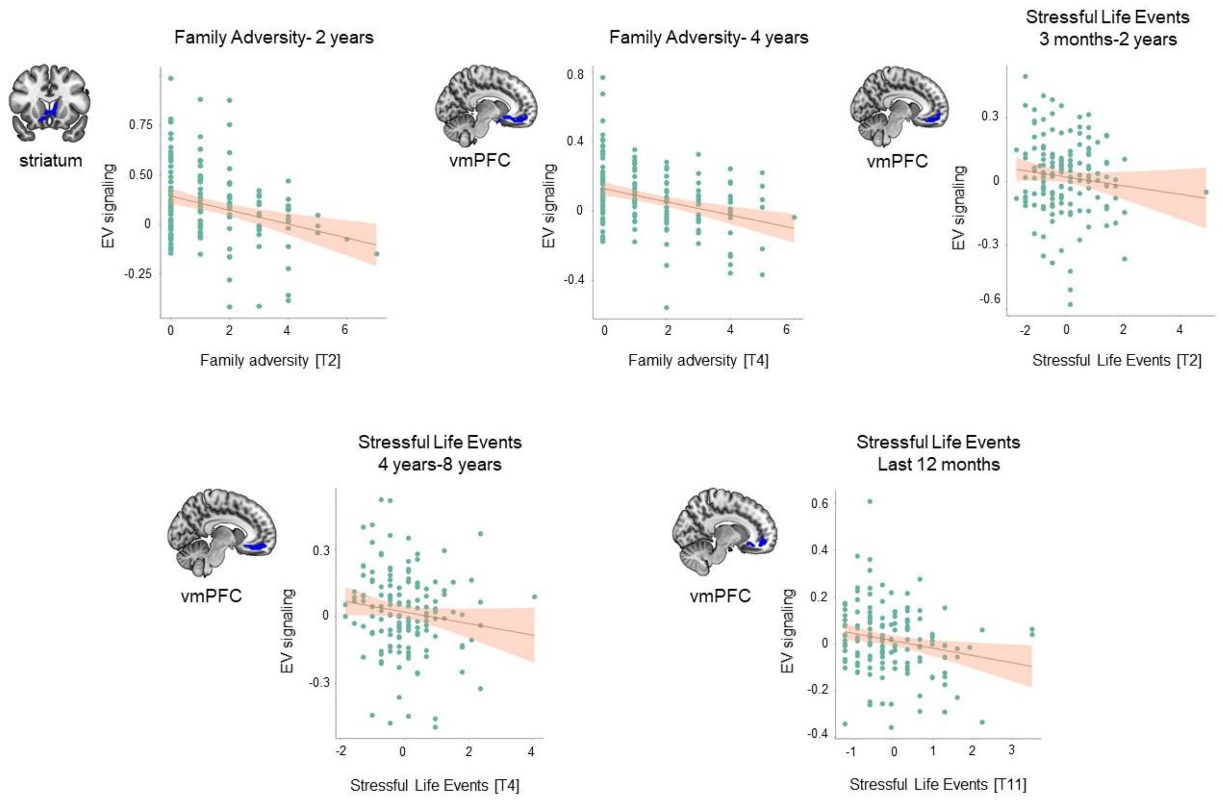

**Figure S4.** Exploratory analyses on the timing effect of adversities. Blue color represents lower expected value signaling in the regions-of-interest. Both measurement time and time-span covering the occurrence of stressful events were reported for stressful life events measurement. Abbreviations: EV, expected values; vmPFC, ventromedial prefrontal cortex.

#### S8. Brain-Behavior Association

The first adversity factor was associated with lower EV encoding in the vmPFC. Lower EV encoding in the vmPFC was further associated with higher internalizing ($r=-0.22$, $p=0.006$), externalizing symptoms ($r=-0.16$, $p = 0.048$). Lower EV encoding in the vmPFC was also related to higher avoidant personality ($r=-0.16$, $p = 0.04$) and ADHD ($r=-0.20$, $p = 0.01$) symptoms (Figure S5). However, these correlations did not survive after the Bonferroni correction.

The third adversity factor was associated with lower EV signaling in the cluster containing the vmPFC, right striatum and pregenual ACC. We did not identify any association between the EV signaling in the large cluster and psychopathology. However, when we examined the effect of specific regions, we found that lower EV signaling in the vmPFC was associated with higher internalizing problems ($r=-0.28$, $p = 0.000$) and externalizing problems ($r=-0.22$, $p =0.006$). In addition, we found a negative correlation between EV signaling in the vmPFC and several ASR subscales including depression($r=-0.24$, $p = 0.003$), avoidant personality ($r=-0.17$, $p =0.03$), somatic problems ($r=-0.21$, $p =0.01$), antisocial personality ($r=-0.18$, $p = 0.02$) and ADHD ($r=-0.24$, $p = 0.002$) subscales (Figure S6). However, only the correlations with depression and ADHD subscales survived after the Bonferroni correction.

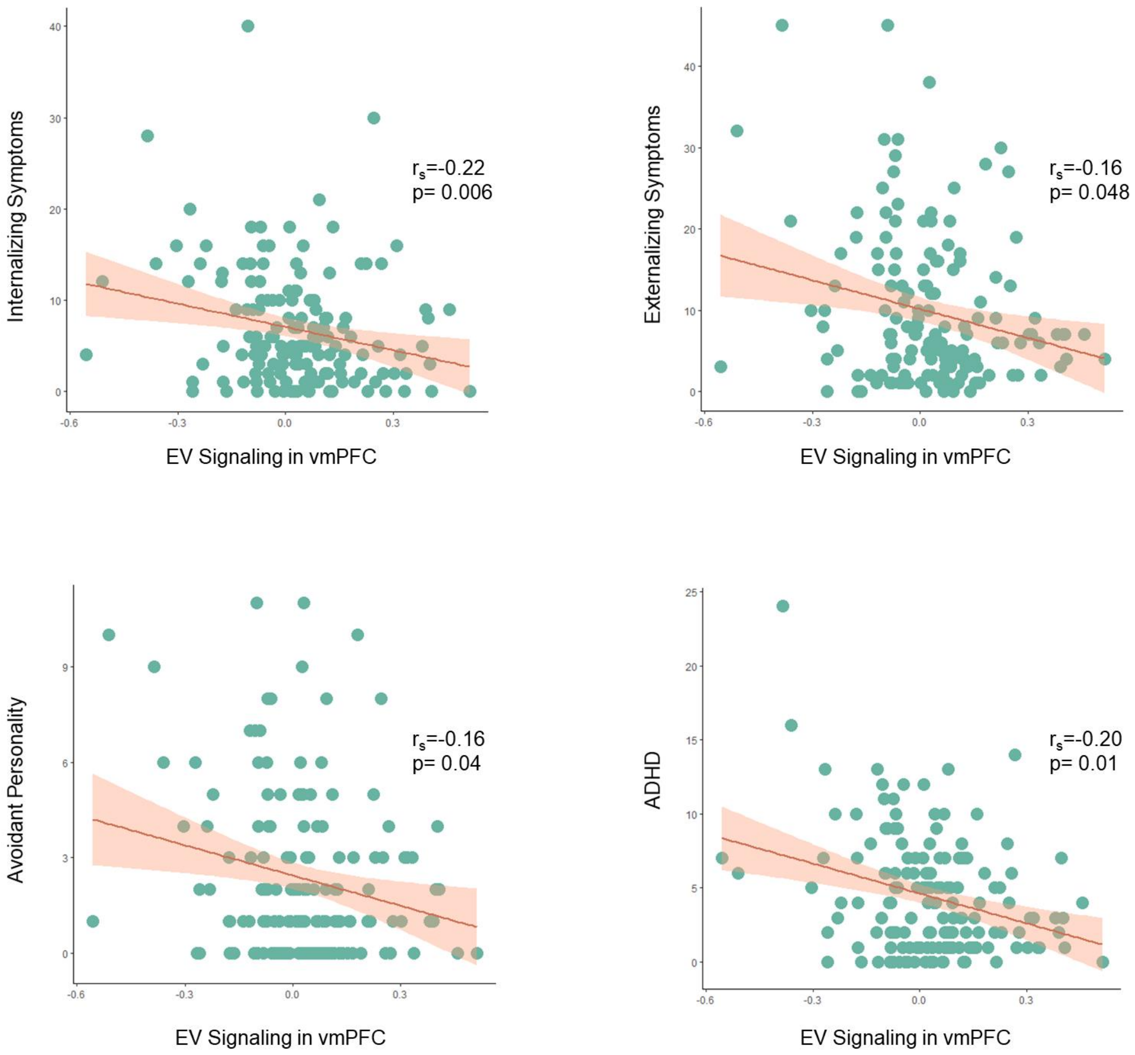

**Figure S5.** Adversity Factor 1: The associations between expected value signaling in ventromedial prefrontal cortex and psychopathology symptoms.

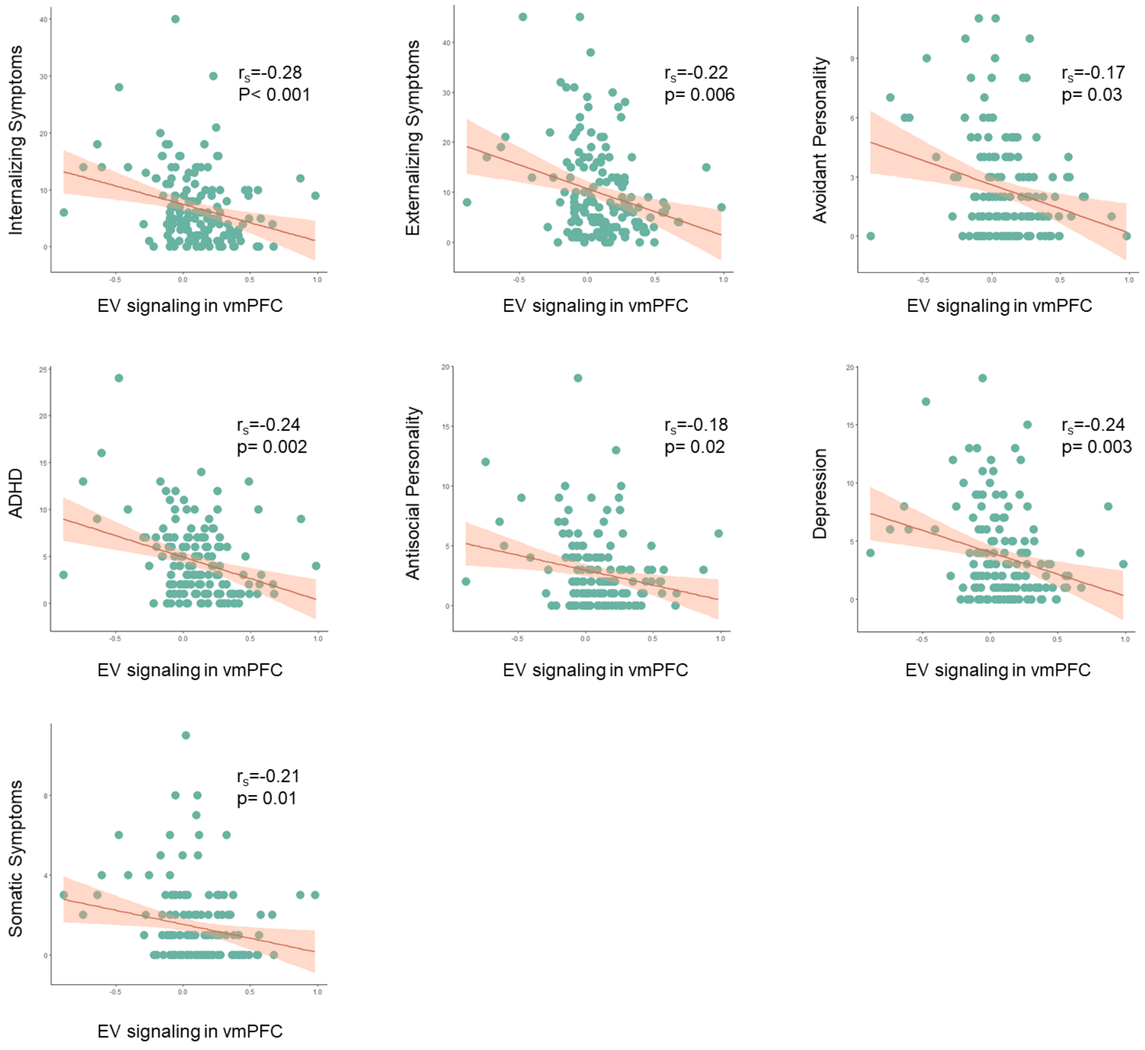

**Figure S6.** Adversity Factor 3: The associations between expected value signaling in ventromedial prefrontal cortex and psychopathology symptoms.
